# A data-driven analysis of spatiotemporal cues and experience accumulation effects for pitch type prediction

**DOI:** 10.1101/2025.10.29.685437

**Authors:** Ryota Takamido, Chiharu Suzuki, Hiroki Nakamoto

**Author notes:** Corresponding author: (RT).

## Abstract

Conventional sports anticipation studies have primarily relied on hypothesis-testing paradigms that target predetermined cues. However, such approaches risk overlooking unanticipated sources of predictive information. This study addresses this limitation by introducing data-driven analysis using machine learning (ML) models as a complementary approach to conventional experimental research. Given that predictive cues embedded within movements can enhance the prediction accuracy of ML models, the proposed analysis identified spatiotemporal cues for prediction and quantified the effects of accumulating opponent-specific information across trials. Motion-capture data were collected from eight collegiate baseball pitchers, and joint-angle time series were analyzed using logistic regression models to predict pitch type (fastball vs. breaking ball). Specifically, two analyses were conducted: (1) a sliding time-window analysis to identify when and where predictive cues emerged within target motions and (2) a set-size analysis to evaluate how prediction accuracy varied with dataset size. The main results revealed that (1) predictive cues were distributed across the entire body, but models integrating whole-body information achieved the highest accuracy; (2) informative cues emerged in most body regions around the initiation of the pitcher’s weight shift; (3) the accumulation of opponent-specific information had a pronounced effect up to approximately 30 pitches; and (4) substantial individual differences existed in when and which cues were effective for pitch-type prediction. Although alignment with human athletes must be carefully examined in future work, these findings highlight the utility of the proposed analysis as a complementary approach to conventional hypothesis-testing experiments.

## 1. Introduction

Anticipation—defined as the cognitive ability to observe others’ movements, infer their intentions and goals, and predict forthcoming events—is a critical factor in athletic performance in dynamic, high-speed sports [1]. Given that the combined time required for perception, decision-making, and motor execution often exceeds the time available to athletes (e.g., the ball’s flight time in hitting), anticipating future events and initiating actions in advance are essential for achieving task goals [2]. Therefore, understanding how skilled athletes decode information from others, infer their intentions, and generate accurate predictions remains a key topic in sports science.

From a methodological perspective, numerous studies have investigated experts’ anticipation skills using controlled experimental paradigms designed for hypothesis testing. In these studies, representative visual stimuli—such as an opponent’s pitching motion or early ball-flight information in baseball hitting [3,4]—were presented to participants under specific experimental manipulations (e.g., occluding a particular body part). Participants’ verbal or motor responses were collected and statistically analyzed to test the relevant hypotheses. Consequently, several unique characteristics underlying experts’ superior anticipation skills have been identified, including their ability to utilize temporally advanced kinematic information from opponents [5,6], spatially localized cues from specific body parts [7,8], and individual differences associated with anticipatory ability [9,10].

However, a fundamental limitation of conventional sports anticipation research is its lack of data- driven and exploratory approaches, as most studies have focused heavily on hypothesis-testing experimental designs. Although hypothesis-testing experimental studies rigorously verify knowledge derived from experienced coaches and athletes, they are less likely to yield insights beyond the experimenter’s prior assumptions [11,12]. Most previous sports anticipation studies selected candidate cues presumed to contribute to predictions and manipulated them experimentally. Consequently, the main findings have largely been restricted to confirming or rejecting proposed hypotheses and their associated analytical results. If accurate prediction is achieved with only a limited number of spatiotemporal cues, participants may not need to use all available information, creating a risk that cues not commonly used across the studied group will be overlooked (Fig 1a).

**Fig 1.**
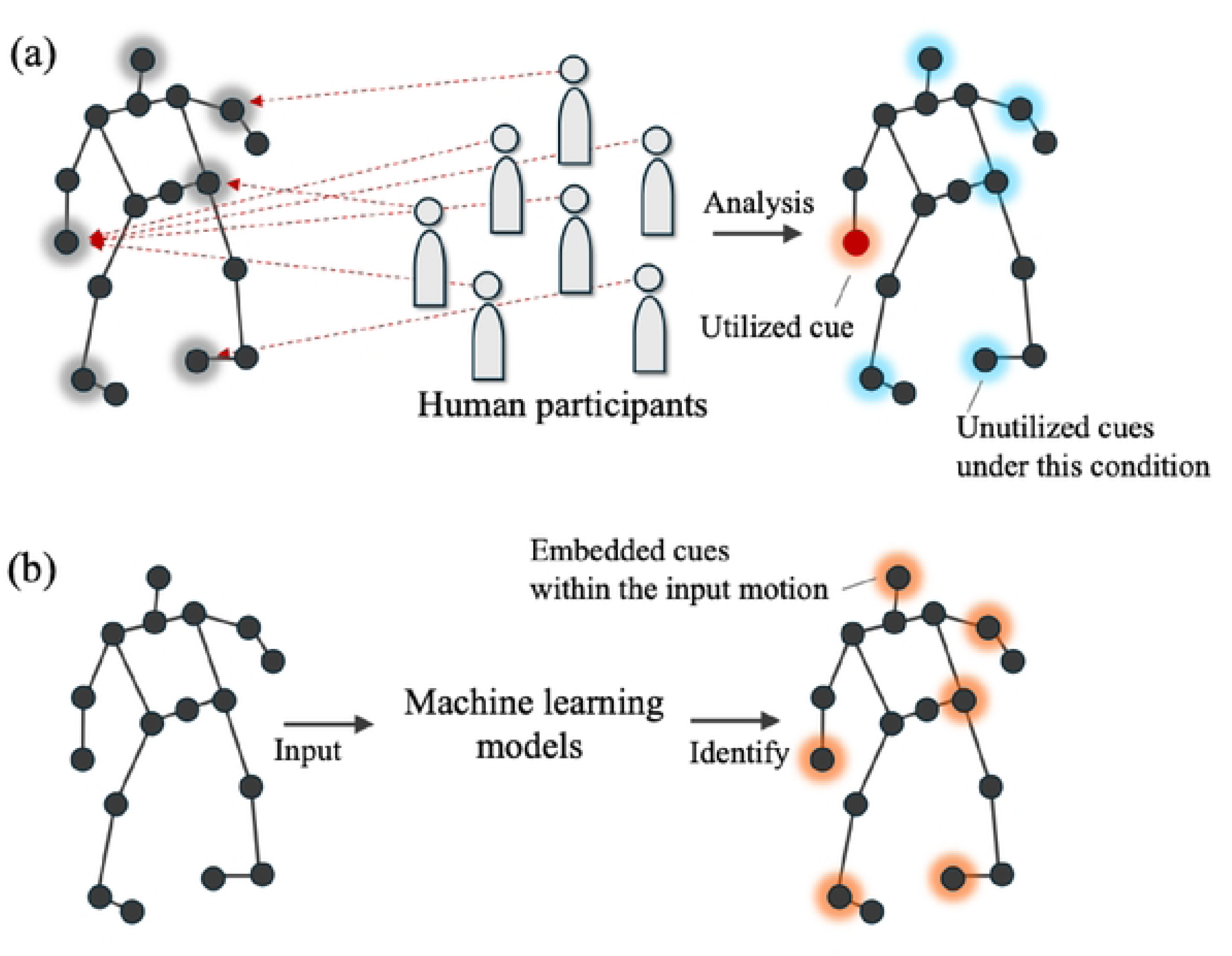
A schematic comparison between (a) a hypothesis-testing experimental paradigm and (b) a data-driven machine learning analysis.

Achieving a comprehensive understanding of athletes’ anticipatory skills requires considering two key aspects: (1) the types of spatiotemporal prediction cues potentially embedded in an opponent’s movement and (2) which of those cues athletes selectively use. This distinction enables researchers to determine whether the differences in prediction cues across studies arise from variations in the information embedded within the visual stimuli or from the characteristics of the participants in each study. However, prior studies have predominantly focused on identifying cues that athletes explicitly use. A fully comprehensive method for systematically examining when and where, within an opponent’s full-body motion sequence, the cues that enhance predictive accuracy are embedded has not yet been established. Although several previous studies [13,14] have employed bottom-up approaches, such as principal component analysis, to identify prediction cues, these methods compute weights across the entire time sequence. As a result, these approaches enable the identification of spatial cues but make it difficult to determine temporal cues directly.

Addressing the shortcomings of conventional methods, data-driven analyses based on machine learning (ML) models have recently gained attention as a complementary approach [12,15,16]. These studies analyzed measured data using ML models to identify new insights, models, and patterns (e.g., [17,18]). Specifically, in the context of sports anticipation, we argue that ML models can help to identify informative cues that enhance athletes’ predictive skills and the effects of their accumulation across trials (Fig 1b). From an information-theoretic perspective, the information embedded in an opponent’s movements should, in principle, be quantifiable directly from motion data by assessing how its use improves prediction accuracy—that is, reduces uncertainty. Similarly, the accumulation of opponent-specific information across trials should be quantifiable by evaluating how much it promotes the separation of informative cues from noise. Although existing anticipation studies have rarely focused on information accumulation across trials because of practical constraints with human participants (e.g., limits on the number of trials), the ability to accumulate opponent-specific information and refine predictions over time may be a crucial factor in real-game situations. These data-driven insights can deeper understanding of sports anticipation skills by complementing conventional hypothesis-driven experiments.

Building on this background, this study aims to develop a novel ML-based analysis that provides data-driven insights into athletes’ anticipation skills. Specifically, we assumed that motion information improving ML prediction accuracy—that is, reducing uncertainty—can be regarded as a potential cue contributing to anticipation skills. Building on this background, we conducted two analyses: (1) a data-driven identification of spatiotemporal cues that enhance sports anticipation skills and (2) a quantitative evaluation of how the accumulating opponent-specific information across trials influences prediction accuracy. Although some recent studies have applied ML models to predict sports movement outcomes [19–22], these approaches have several limitations, including limited explainability [21], restriction of identified cues to spatial (body-part) information [22], and reliance on non-athlete motion data [19,20]. Unlike previous approaches, our method incorporates athletes’ motion data and identifies interpretable spatiotemporal cues while further providing a novel analysis of how accumulating opponent-specific information across trials contributes to improved anticipation skills.

## 2. Materials and methods

In developing the proposed analysis, this study selected pitch-type prediction in baseball (fastball vs. breaking ball) as the target task, given the complexity of the motion information provided by the opponent (i.e., the pitcher) and the importance of accurate prediction for coping with strict temporal constraints during real-game striking actions [2]. For the analysis, we constructed individual machine-learning models for each pitcher, analogous to the way participants in experimental studies typically learn predictive models based on the actions of a single actor. This design also reflects real competitive contexts, in which batters adapt to individual pitchers by progressively accumulating information across pitches. The source code and dataset used for the analysis are publicly available on GitHub (https://github.com/takamido/Pitch_type_pred_ML).

### 2.1 Overview of the proposed analysis

This analysis was inspired by recent neuroscience research that aimed to understand the spatiotemporal characteristics of electroencephalography (EEG) signals by validating the decoding accuracy of ML models [23,24]. In this study, we extend this framework to identify key spatiotemporal cues for sports anticipation embedded within complex athletic movements and to evaluate how the accumulation of these cues across trials influences prediction accuracy. Figs 2 and 3 present schematic illustrations of the proposed analysis. Fig 2 depicts a general framework for training and testing a pitch-type classification model using ML, and Fig 3 presents a more specific analysis design tailored to the context of sports anticipation.

**Fig 2.**
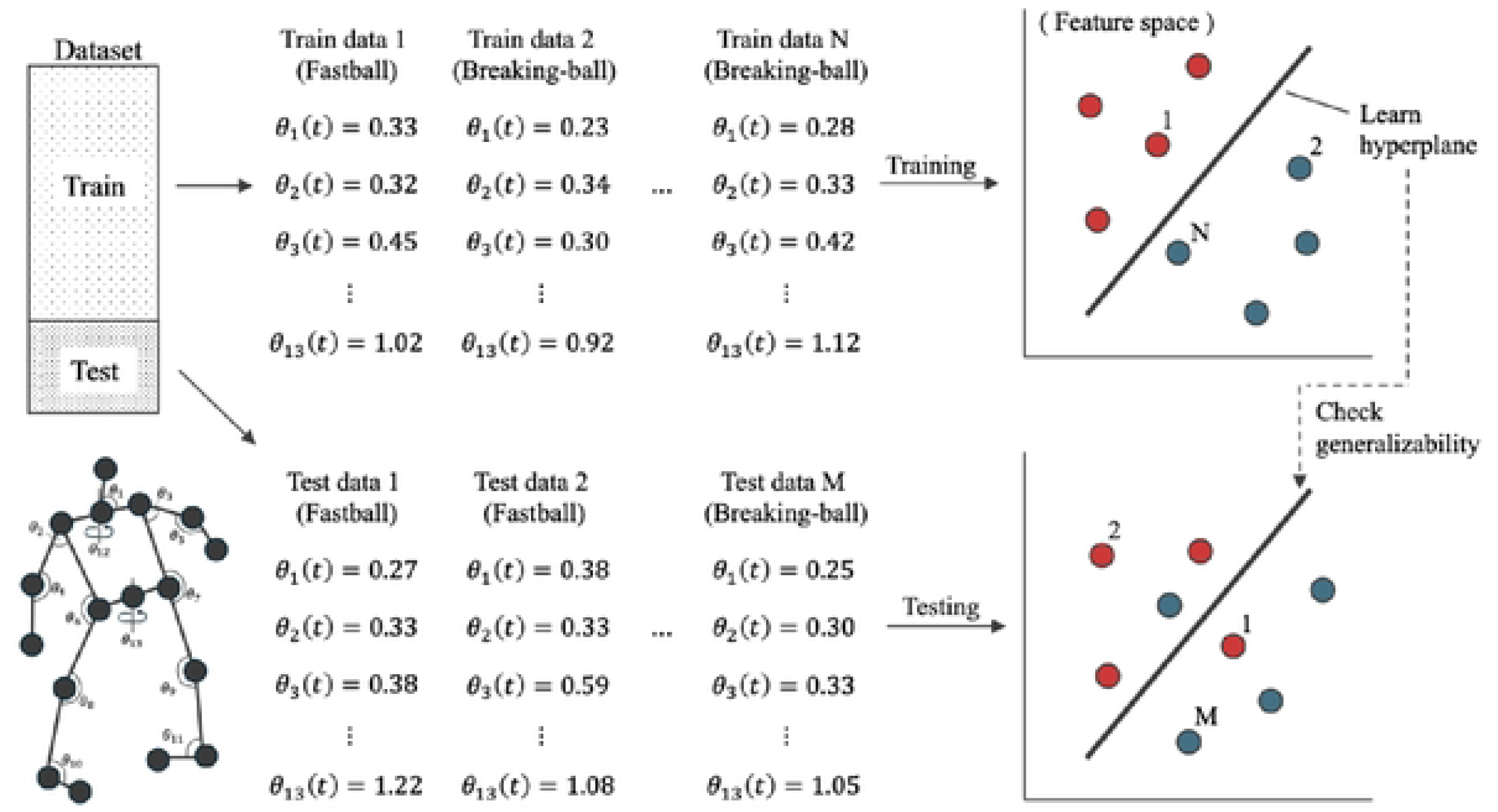
Schematic image of ML-based pitch-type prediction using joint angles at time *t* as features.

**Fig 3.**
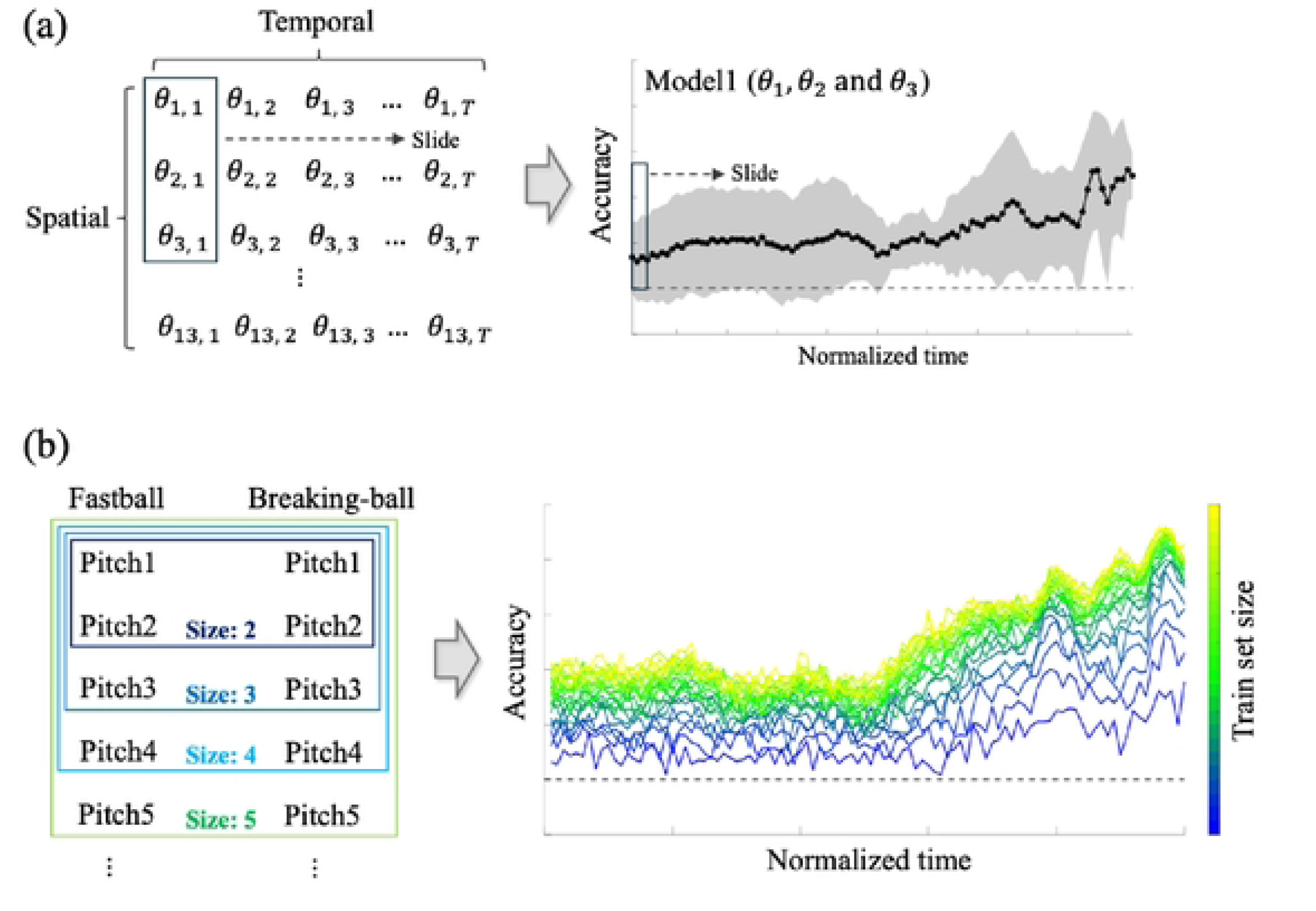
Schematic illustration of the two analyses conducted in this study. (a) Analysis 1: Sliding time-window analysis for identifying spatiotemporally informative cues. (b) Analysis 2: Set-size analysis for evaluating how the accumulation of information across trials influences the prediction accuracy of the ML model.

Conceptually, the task of the ML model is to decode the spatiotemporal motion data—defined in this study as a time series of joint angles—by leveraging the information contained in the training data to determine whether it corresponds to a fastball or a breaking ball (Fig 2). If the model achieves high prediction accuracy using the given information, it can be inferred that the corresponding spatiotemporal features may contribute to improved anticipation performance.

Building on this property of ML, two analyses were conducted in this study. In Analysis 1, we used data-driven ML analysis to reveal when and where the information for pitch-type classification was embedded within the spatiotemporal motion data (Fig 3a). In this analysis, we repeatedly trained the ML model using sliding time windows and evaluated its prediction accuracy, enabling continuous characterization of temporal changes in prediction probability. Furthermore, restricting the input to specific joints (e.g., those in the throwing arm) allowed identification of the joints and time points at which their information contributed to improving the prediction accuracy of the ML model. The prediction results obtained from the individual pitcher models were aggregated and statistically analyzed to verify the presence of spatiotemporal features shared across individuals.

Furthermore, in Analysis 2, we evaluated how the prediction accuracy changes over longer timescales as information about the target pitcher accumulated across trials, mirroring how batters incrementally learn about their opponents during real matches. This evaluation was achieved by varying the size of the training dataset (Fig 3b). Through random resampling of the training data and repeated training and testing with different dataset sizes, ML enables data-driven evaluation of the incremental contribution of each additional trial and its impact on the prediction accuracy.

Although the transferability of these analytical results to human athletes should be interpreted with caution, this novel analysis is expected to complement conventional hypothesis-testing experiments and serve as a foundation for follow-up studies that advance understanding of predictive abilities.

### 2.2 Dataset collection and processing

The dataset consists of data from eight college league pitchers (174.1±4.1 cm, 74.91±3.8 kg, 19.87 ± 1.0 years). All pitchers had more than 10 years of playing experience. This study used previously collected, unpublished data. An opt-out informed consent framework was adopted in accordance with the university’s ethical guidelines. Information about the study, including its purpose and data usage, was made publicly available to ensure transparency. Individuals whose data were included had a clear opportunity to decline participation. The requirement for written informed consent was waived by the Institutional Ethics Committee of the National Institute of Fitness and Sports in Kanoya, which approved the study procedure (approval number: 25-1-26). The data were accessed for research purposes between September 1, 2025, and October 27, 2025, during the implementation of this study.

All data were fully anonymized prior to analysis, and no identifying information was available to the authors. All procedures adhered to the principles of the Declaration of Helsinki.

For dataset collection, the motion of each pitcher was measured at a sampling rate of 200 Hz using 16 synchronized optical motion-capture cameras (Raptor-E and 16 Kestrel2200 cameras, Motion Analysis Corp, Santa Rosa, USA). Pitchers threw random pitch types at random locations during the experiment to simulate real game-like conditions. The positions of 15 joints—the parietalis (head), both sides of the acromions (shoulders), lateral epicondyles of the humerus (elbows), radial styloid processes (wrists), greater trochanters of the femur (hips), lateral condyles of the femur (knees), and heels and tops of the shoes (toes)—were extracted from the measured data. Based on the measured joint-position data, time-series features were computed as follows: ten three-dimensional joint angles—both shoulders (𝜃_2_, 𝜃_3_), elbows (𝜃_4_, 𝜃_5_), hips 𝜃_6_, 𝜃_7_, knees 𝜃_8_, 𝜃_9_, and ankles (𝜃_10_, 𝜃_11_)— as illustrated in Fig 2, and three two-dimensional angles (sagittal-plane angle between the head and shoulder vector and the horizontal axis of the ground (𝜃_1_) and the horizontal-plane rotation angle of the shoulder (𝜃_12_) and hip (𝜃_13_)). Data from the left-handed pitchers were mirrored to standardize the meaning of each angle. Subsequently, for each pitch, features from the onset of the pitcher’s wind-up to the point of ball release were extracted and normalized to 101 time points. Wind-up onset was defined as the frame at which the vertical velocity of the leading knee dropped below 5% of its maximum value, traced backward from the moment when the knee reached its peak vertical position. The ball release was defined as the frame at which the resultant velocity of the throwing-arm wrist reached its maximum. Illustrations of the 20% intervals of the normalized motion are shown in Fig 4. The pitching motion data for each pitcher are available at GitHub (https://github.com/takamido/Pitch_type_pred_ML).

**Fig 4.**
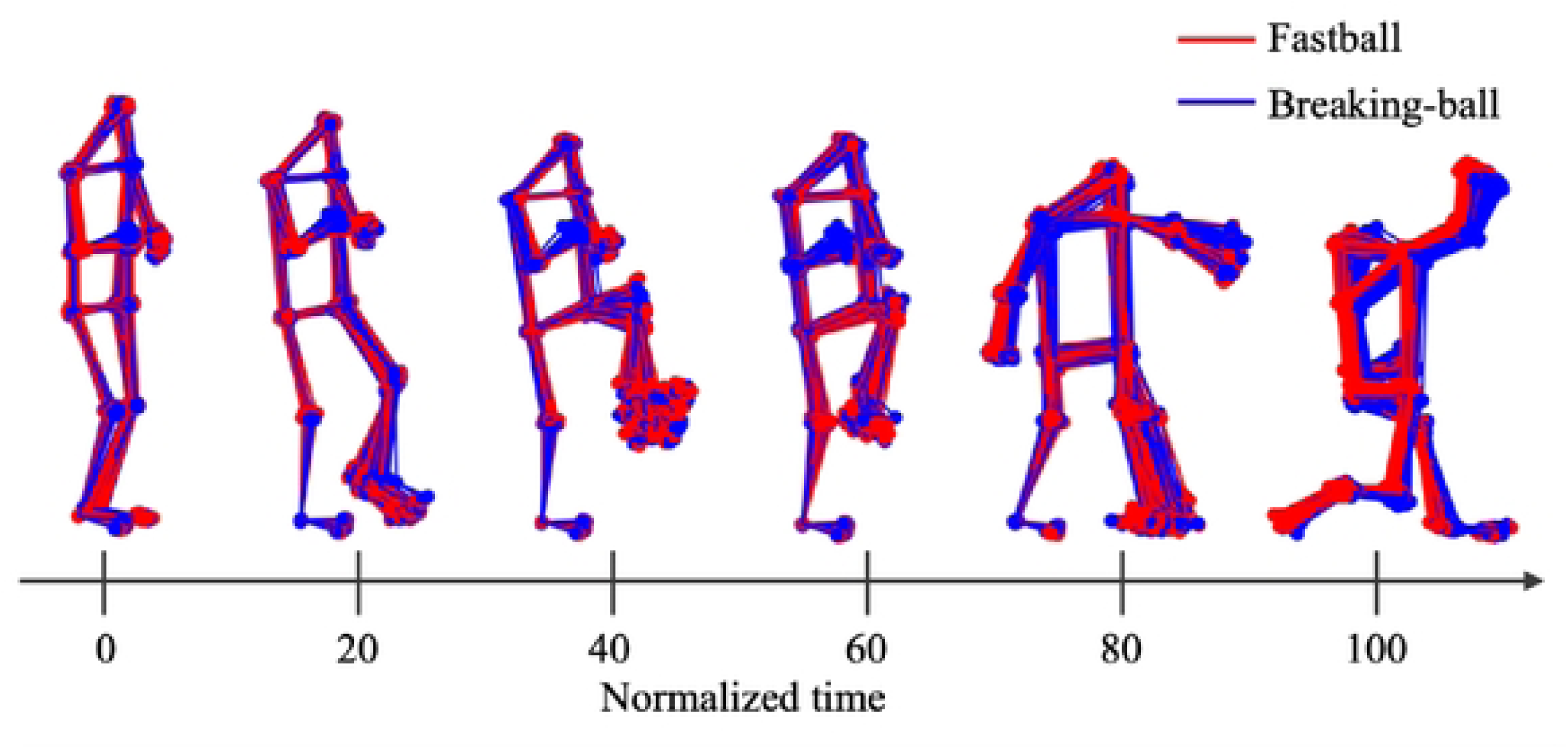
Illustrations at 20% intervals of the normalized motion, using Sub1 as an example.

The ball velocity (initial speed) of each pitch was also recorded as a performance index using the TrackMan system (TrackMan). Pitch type information was collected based on the participants’ self-reports. A pitch was classified as a breaking ball if its average velocity was at least 10 km/h (6.21 mph) lower than the average fastball velocity of the pitcher. The breaking balls include change-ups, curveballs, sliders, sinkers, or splitters. Table 1 presents the average velocities of fastballs and breaking balls, along with the total number of pitches of each type for each pitcher. It should be noted that because the pitchers simulated game-like situations by throwing random pitch types, the distribution of pitches across types was unbalanced.

**Table 1.** Summary of the velocities and pitch counts of fastballs and breaking balls for each pitcher.

|  | Fastball pitch | Breaking-ball pitch | Mean fastball | Mean breaking-ball |
| --- | --- | --- | --- | --- |
|  | number | number | velocity (mph) | velocity (mph) |
| Sub1 | 56 | 62 | 69.44 | 62.83 |
| Sub2 | 57 | 48 | 76.09 | 63.74 |
| Sub3 | 37 | 48 | 76.41 | 64.65 |
| Sub4 | 55 | 59 | 73.83 | 66.16 |
| Sub5 | 56 | 44 | 75.79 | 67.1 |
| Sub6 | 47 | 50 | 80.18 | 65.55 |
| Sub7 | 30 | 40 | 75.41 | 63.40 |
| Sub8 | 80 | 31 | 76.17 | 63.86 |

### 2.3 ML model for the analysis

Consistent with previous approaches used for EEG data analysis [23], logistic regression was employed as the classification model. Logistic regression is a linear method that estimates the probability of class membership using a logistic function applied to the weighted sum of input features.

Specifically, given the joint angles at time *t* as input (𝚯_𝐭_), the logistic regression model calculated the probability that the pitch was a fastball using the following equation:

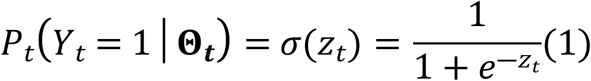

where 𝑧_𝑡_ = 𝛽_0_ + 𝛽_1_𝜃_1,𝑡_ + 𝛽_2_𝜃_2,𝑡_ +… + 𝛽_13_𝜃_13,𝑡_ represents the linear combination of input features, and 𝜎 is the sigmoid function. When prediction is performed using only specific joint angles, for example 𝜃_1_ and 𝜃_2_, the expression reduced to 𝑧_𝑡_′ = 𝛽_0_ + 𝛽_1_𝜃_1,𝑡_ + 𝛽_2_𝜃_2,𝑡_.

Based on Equation (1), the logistic regression model sought to determine a separating hyperplane between the fastball and breaking-ball data points up to a 13-dimensional joint-angle space (Fig 2). Intuitively, this process corresponds to classifying the pitch type according to the relative configuration of joint angles, for example, interpreting a motion with a deeply flexed lead knee and an extended throwing shoulder as a fastball. Postural information is a common predictive cue that has been identified in many previous studies [25]. When multiple joint angles are available as input, those that contribute more to prediction are assigned larger weights (𝛽). Although more complex classification models could be employed, we used logistic regression in this study because it relies on relatively simple information, provides high interpretability and explainability, and allows the identified information to be potentially useful for human athletes.

### 2.4 Analysis 1: sliding time window analysis

In Analysis 1, we aimed to identify the spatiotemporal features relevant to pitch-type prediction by applying a sliding time-window approach to the joint-angle time series. Specifically, for each pitcher, the logistic regression model was trained and tested using the feature vectors corresponding to a given time point *t* within the 101 normalized time points. We continuously evaluated classification performance across the entire pitching motion by sliding the time window and repeatedly training and testing the model. We further examine spatial importance by conducting independent evaluations under seven conditions: one using all 13 joint features simultaneously and six dividing the joints according to body regions (e.g., throwing arm: 𝜃_2_ and 𝜃_4_, leading arm: 𝜃_3_ and 𝜃_5_, trunk tilt: 𝜃_1_, 𝜃_6_ and 𝜃_7_, trunk horizontal rotation: 𝜃_12_ and 𝜃_13_, pivot leg: 𝜃_8_ and 𝜃_10_, leading leg: 𝜃_9_ and 𝜃_11_). We repeated the following procedure 100 times for each time point *t*, to ensure robust estimation, resulting in 7 × 101 × 100 = 70,700 evaluations per pitcher:

1. Data partitioning: For each pitch type, 25 trials ( 50 in total) were randomly sampled as training data, and five trials ( 10 in total) were sampled as test data. This procedure corrected data imbalances across pitchers and maintained a consistent chance level. In addition, this balanced sampling procedure ensured that the classifier was trained with equal numbers of fastball and breaking-ball trials while keeping the test sets independent. Similar to bootstrap methods [26], this approach reduced the sensitivity to extreme data points by repeatedly utilizing subsets of the overall dataset, thereby stabilizing the mean values used in subsequent statistical analyses.
2. Preprocessing: Input features were standardized to a standard normal distribution based on the training set. Normalization parameters (mean and variance) estimated from the training data were then applied to normalize the test data. This procedure prevented information leakage, which would otherwise have occurred if the training and test data were normalized together.
3. Model training and evaluation: A logistic regression model with an iterative solver (maximum 1,000 iterations) was fitted to the training set. The *lbfgs* solver implemented in the *sklearn.linear_model* library in Python was employed with L2 regularization. After fitting, classification was performed on the test data, and the prediction accuracy of the independent test set was recorded.

As a result of the above analysis, the average accuracies at 101 time points were obtained for each of the eight pitchers under the seven input feature conditions. Based on these data, we conducted the following statistical analyses to identify spatiotemporal information that exhibited statistically significant discriminative accuracy. First, to examine whether the logistic regression model could predict pitch type from full joint-angle information at a level significantly above chance (0.50), we applied a cluster-based permutation test [27] using the mean and standard deviation across the eight pitchers under the all-joint condition. Cluster-based permutation testing was conducted using 10,000 permutations. The cluster-forming threshold was set at *p* < 0.05 (one-sided), and the cluster-level significance criterion was set at *α* = 0.05. Cluster mass was defined as the sum of *t*-values across contiguous time points above the threshold, and significance was determined relative to the permutation-based null distribution generated through random sign flips. Clusters shorter than ten consecutive time points (corresponding to approximately 0.2 s) were excluded from the analysis.

Furthermore, for each of the six individual body-region conditions, a cluster-based permutation test was applied to the averaged data across the eight pitchers using the same procedure. The initial cluster-forming significance criterion was set at α = 0.05. We accounted for multiple comparisons by adjusting the significance levels of the identified clusters using the Holm–Bonferroni method across the six body regions.

### 2.5 Analysis 2: set size analysis

In Analysis 2, we evaluated how the prediction accuracy of the ML model changed over longer timescales as information about the target athlete accumulated across the trials. Specifically, for each pitcher, the training dataset size was incrementally increased from 2 to 25 trials per pitch type (i.e., 4– 50 trials in total), and the logistic regression model was repeatedly trained and tested. The test dataset was fixed at five trials per pitch type (ten trials in total), independent of the training dataset size. For each dataset size, the training and test data were randomly shuffled, and the accuracy was evaluated 100 times at each of the 101 time points. Data preprocessing and logistic regression model fitting were performed using the same procedures as in Analysis 1.

From the above analysis, the mean prediction accuracies were obtained for each pitcher, and the dataset size was calculated at each of the 101 time points. Based on these values, we computed the mean accuracies for dataset sizes of 2, 5, 10, 15, 20, and 25 trials per pitch type. Differences in the mean accuracy across the five intervals (2–5, 5–10, 10–15, 15–20, and 20–25 trials) were then calculated, and the means and standard deviations were obtained across individuals. We reduced the impact of multiple comparisons and mitigated noise by making comparisons over larger spans rather than computing differences at single-trial increments. We verified whether the increases in accuracy across each interval were significantly greater than the chance level (0.0) by applying cluster-based permutation tests to all five intervals. These tests were conducted following the same procedure as in Analysis 1, and significance levels were adjusted for multiple comparisons using the Holm–Bonferroni method.

## 3. Results

### 3.1 Analysis 1: sliding time window analysis

Fig 5 illustrates the mean prediction accuracy and standard deviation under the all-joint feature condition at each time point for each of the eight pitchers. Fig 6 summarizes the overall mean and standard deviation across all pitchers. As shown in the figures, although individual differences existed in overall accuracy and temporal patterns, the ML model demonstrated prediction accuracy above the chance level across most time points when all joint features were used, showing a general trend of increasing accuracy as the ball release approached. The cluster-based permutation test of mean accuracy across pitchers under the all-joint condition revealed that the entire interval constituted a single significant cluster (*p* < .01).

**Fig 5.**
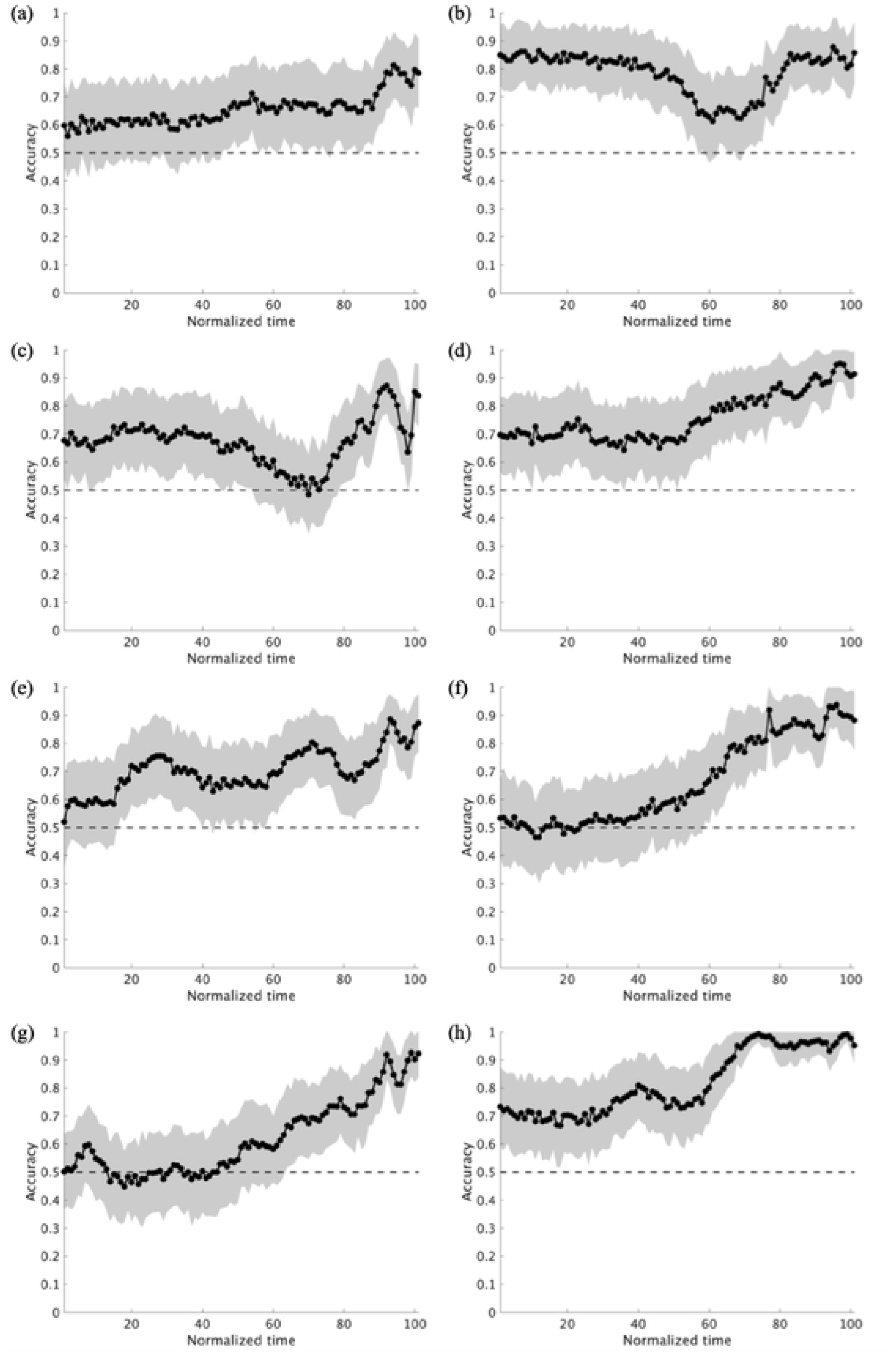
Mean prediction accuracy and standard deviation for each pitcher at each time point using full-joint information. Panels (a)–(h) show the individual plots for subjects 1–8.

**Fig 6.**
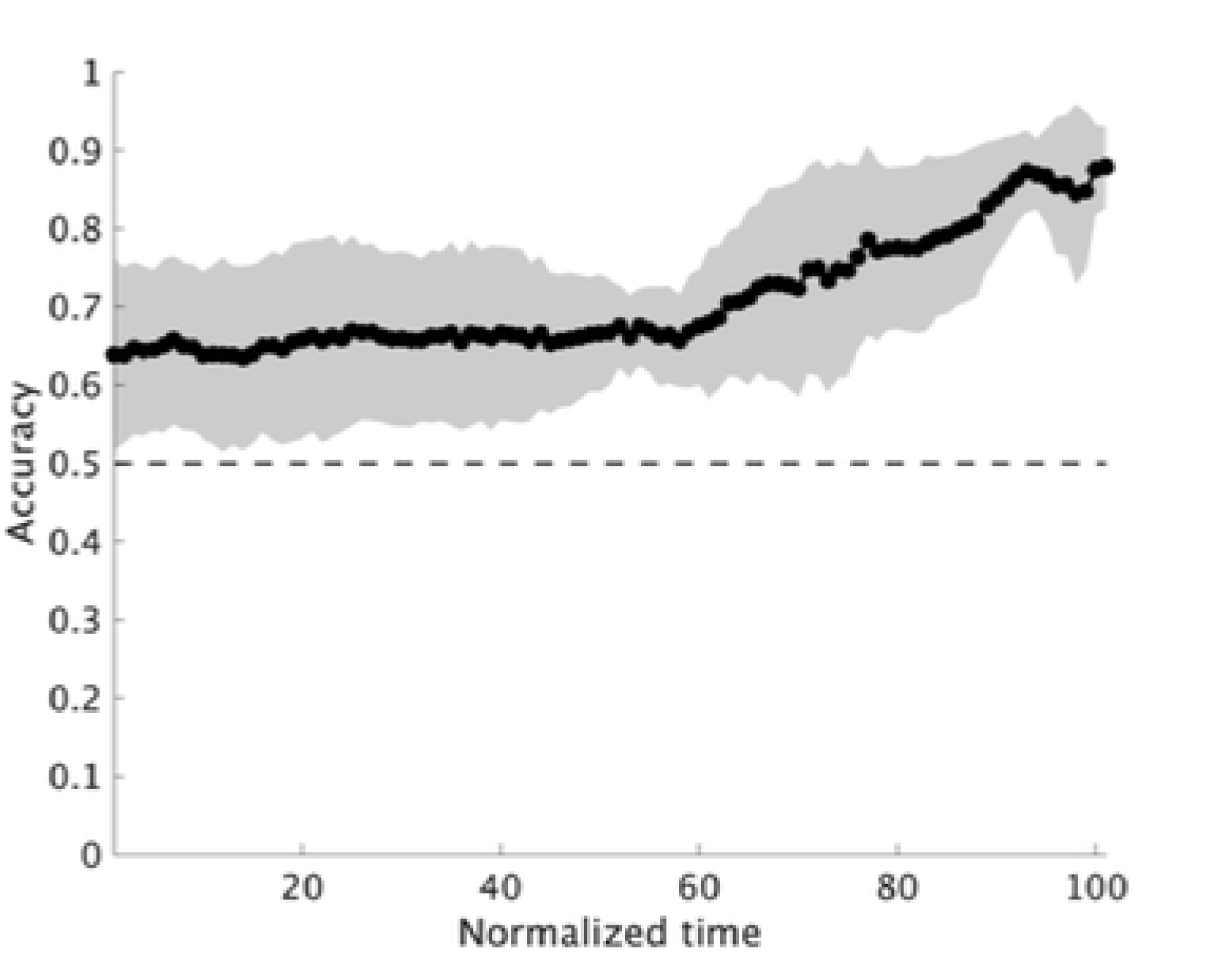
Overall mean prediction accuracy and standard deviation across the eight pitchers using full-joint information.

Fig 7 plots the mean prediction accuracy for each of the six body regions, with results of all eight pitchers overlaid. The cluster-based permutation test for six regions revealed that classification based on the throwing-arm (8%–100%), leading arm (60%–100%), trunk tilt (0%–58% and 65%–100%), trunk horizontal rotation (66%–100%), pivot leg (65%–100%) and leading leg (70%–96%) achieved prediction accuracies significantly above the chance level (*adjusted p* = 0.019, <.01, 0.016, 0.013, <.01, <.01 and <.01, respectively). Although a more detailed discussion of the identified spatiotemporal information is provided later, these results suggest that (1) all regions contain significant information for discriminating pitch types by the ML model, (2) some information enhances prediction accuracy even at early time points, and (3) the final prediction accuracy of the ML model at 100% (release) using information from individual regions (Fig 8) is approximately 15%–20% lower than when using information from all regions combined (Fig 6).

**Fig 7.**
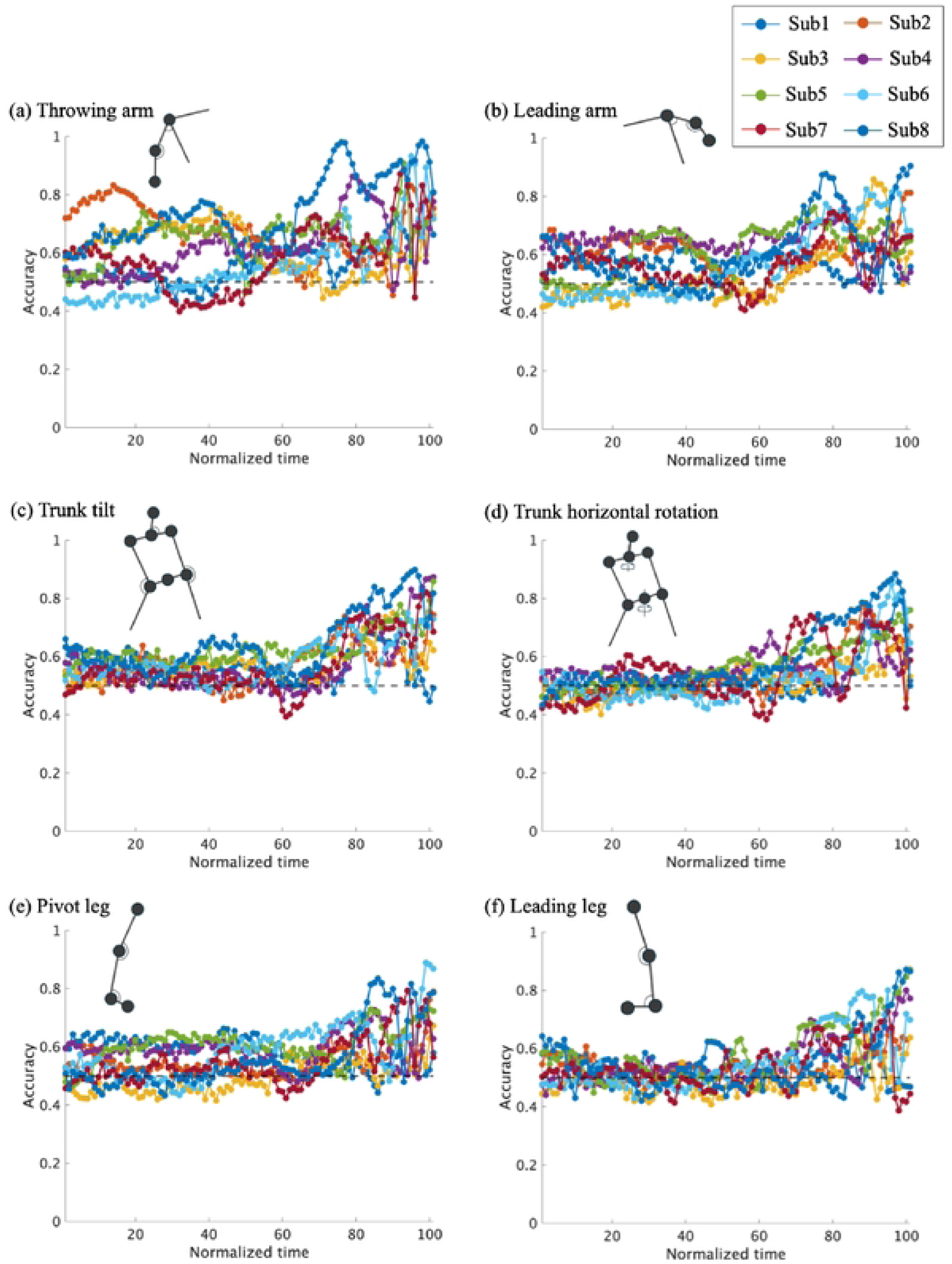
Mean prediction accuracy for each of the six body regions, with results from all eight pitchers overlaid.

**Fig 8.**
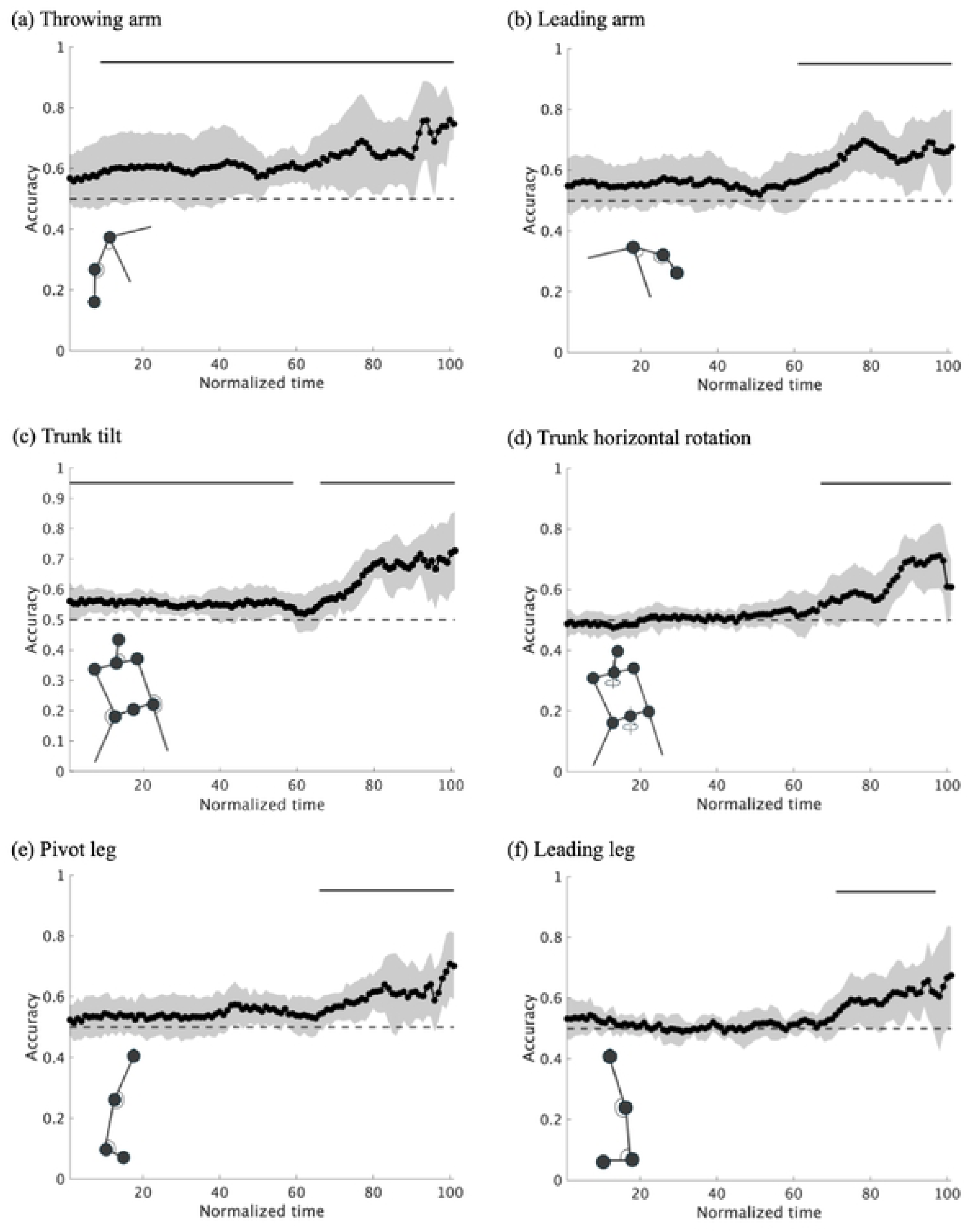
Mean prediction accuracy and standard deviation across eight pitchers for each of the six body regions. Solid lines indicate significant intervals (*p* < .05).

### 3.2 Analysis 2: set size analysis

Fig 9 shows the relationship between dataset size and prediction accuracy at each time point for each pitcher. According to the cluster-based permutation test results, significant improvements were observed in the following intervals: for training dataset sizes of 2–5 trials at 13%–36% and 38%– 100% (*adjusted p* =0.016 and < .01); for 5–10 trials at 3%–17% and 19%–100% (*adjusted p* =0.018 and < .01); for 10–15 trials at 30%–40%, 50%–73%, and 75%–100% (*adjusted p* =0.020, <.01 and <.01, respectively); and for 15–20 trials at 86%–98% (*adjusted p* <.01) (Fig 10). No significant clusters were observed in the trial interval of 20–25 minutes. These results demonstrate that the impact of adding additional training data on prediction accuracy is more pronounced when the dataset size is small. A detailed discussion of these results, including their practical significance, is provided in the following section.

**Fig 9.**
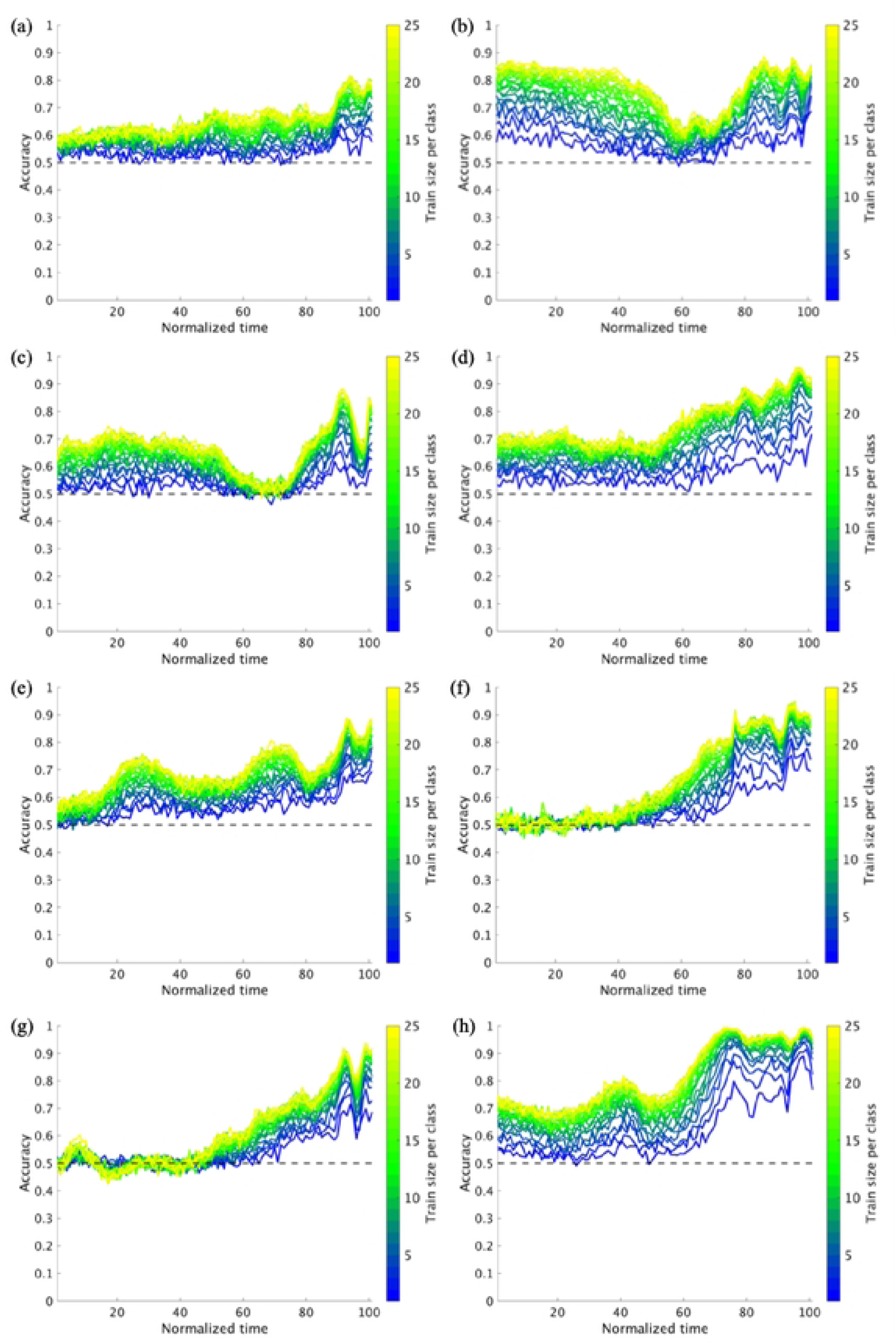
Relationship between dataset size and prediction accuracy at each time point for each pitcher. Panels (a)–(h) show individual plots for subjects 1–8.

**Fig 10.**
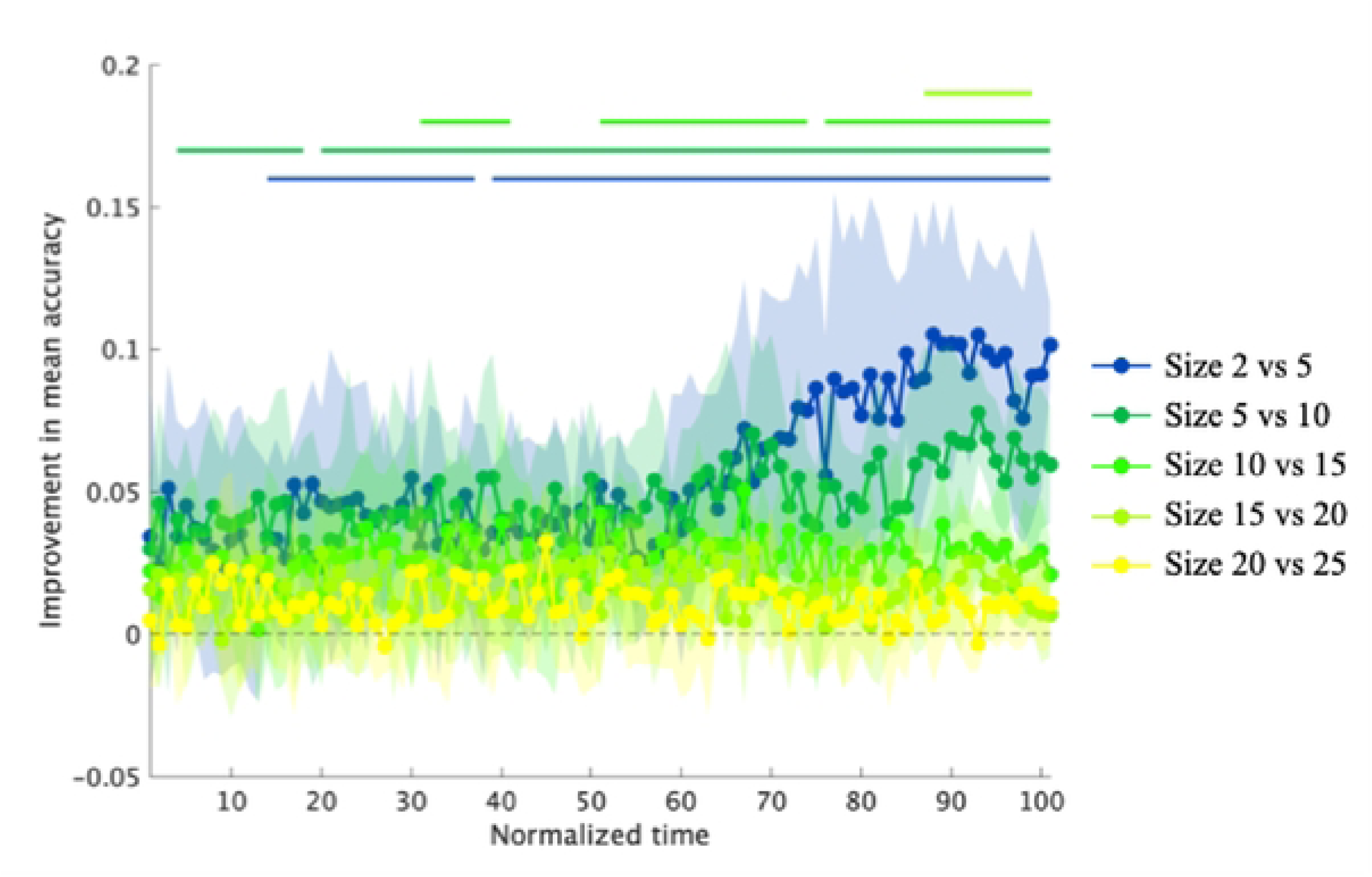
Improvement in mean prediction accuracy with increasing training data. Numbers on the dataset size axis represent the number of trials per pitch type (thus, a size of 5 corresponds to 10 training trials in total). Solid lines indicate intervals where accuracy improvements were statistically significant compared with the chance level (0.0).

## 4. Discussion

First, the results of Analysis 1 indicated that across all six hypothesized body regions, the ML model achieved prediction accuracies above chance level during the 60%–100% interval of the motion sequence (i.e., from around the initiation of the pitcher’s weight shift to ball release). In previous studies, the body parts athletes utilized for prediction—for example, proximal vs. distal [7] or local vs. global cues [8,28,29]—have been important topics. In this context, the results of the proposed analysis suggest that (1) from a purely information-theoretic perspective, all body regions contain information that contributes to prediction accuracy. Therefore, if athletes can access this embedded information, they have the potential to achieve the above-chance prediction accuracy, regardless of cue type (Fig 6). (2) However, models using information from individual regions perform worse than those with access to all joint information (Fig 6 vs. Fig 8); thus, athletes who can integrate information from multiple regions may achieve higher prediction accuracy than those who rely on a single source. Considering that the prediction accuracy at the moment of ball release or ball contact reported in prior studies is approximately 60%–80% [7,29–33], models using a single region were, in terms of accuracy, more comparable to the actual human prediction performance.

Another noteworthy finding of Analysis 1 concerns the temporal aspects of the identified cues. Specifically, the results showed that all body regions exhibited significant clusters from the initiation of the pitcher’s weight shift (approximately 60% of the motion sequence) to ball release (100%). Traditionally, temporal occlusion paradigms for predicting pitch outcomes have been applied near ball release [3,34]. However, these results suggest that cues for pitch-type prediction may emerge at an earlier stage of pitching motion than previously assumed. Moreover, although the difference from chance level was approximately 10%, information from the throwing arm and trunk tilt exhibited significant clusters from the beginning of the wind-up phase (Fig 8). This finding suggests that differences in the intended pitch type are reflected in subtle postural variations during the early phases of the pitching motion. For example, many pitchers alter their grips between breaking balls and fastballs, which may lead to subtle differences in shoulder and elbow joint angles during the wind-up phase. Although it remains unclear whether human athletes recognize such subtle differences, the results of these ML analyses suggest the existence of novel potential cues that may enhance athletes’ hitting performance.

Regarding the results of Analysis 2, with dataset sizes of up to approximately 15 pitches per condition (i.e., 30 in total), significant improvements in prediction accuracy with increasing dataset size were observed across more than 50% of the motion sequences. The greater sensitivity to dataset size in the latter part of the motion is likely due to the increased movement variability near the ball release, with each additional data point contributing more substantially to reducing the uncertainty. From a practical perspective, the improvement in accuracy resulting from the accumulation of pitch information may be associated with the Times Through the Order Penalty, which refers to the phenomenon in which batters tend to improve their performance when facing the same pitcher multiple times. Although the underlying causes of this phenomenon remain under debate [35], the present findings suggest that a reduction in uncertainty and an increase in predictive accuracy resulting from the accumulation of information may be contributing factors. Given that batters can utilize statistical information about their opponent or prior observations made outside the batter’s box, the effect of early information accumulation may be more pronounced.

It is also worthwhile to discuss individual differences in the predictive cues embedded within each pitcher’s motion. As shown in the individual plots (Figs 5, 7, and 9), there were individual differences in terms of the body regions used at different time points and the extent to which such information enabled prediction. For example, while participant 2 achieved an accuracy of approximately 80% using information from the right arm in the early phase, subject 7 demonstrated high accuracy based on information from the trunk in the latter phase. In conventional experimental studies, visual stimuli are typically derived from the movements of a few players; hence, the differences in the cues identified in these studies may be attributable to individual variability. Therefore, it may be ideal to conduct a pre-ML analysis to clarify the spatiotemporal characteristics of the cues embedded in target players before conducting a controlled experiment. Moreover, considering such individual differences among pitchers, skilled batters may adaptively change the spatiotemporal cues they utilize in practical situations.

Finally, this study has some limitations and directions for future work. First, comparing the cues identified using the proposed method with those used by athletes would provide valuable insights. Given that ML models lack the concept of embodiment, such a comparison could further clarify the role of the body and the coupling between perception and action [30,34]in predictive processes. Moreover, because this study focused on intrapersonal training and testing, future studies should examine the generalizability of spatiotemporal cues across individuals. From this perspective, because this study employed a relatively small sample (n = 8), constructing a larger dataset involving more athletes is necessary. In addition, the proposed analysis employed a model that treated postural cues at each time point independently. However, it may be important to consider models specialized for time-series analysis, such as LSTM [36], to better mimic human perception. Finally, as this study focused on pitch type prediction in baseball, future research should examine the applicability of the proposed analysis to a broader range of predictive tasks across sports.

## 5. Conclusion

In conclusion, the data-driven analysis using ML yielded insights that can inform follow-up experiments and deepen the interpretation of existing findings: (1) all body regions contain information that improves the accuracy of pitch type prediction by ML models, but models with access to whole-body information achieve higher accuracy; (2) information from certain body regions significantly enhances prediction accuracy even in the early phase of the pitcher’s motion; (3) prediction accuracy increases substantially as data accumulate up to approximately 15 fastballs and 15 breaking balls; and (4) there are considerable individual differences among pitchers in terms of which information contributes to prediction and at what specific time points. Although alignment with human athletes should be carefully examined in future experiments, these findings highlight the utility of data-driven analysis with ML as a complementary approach to conventional hypothesis-testing experiments. Future research should address topics such as comparing the cues identified by the model with those utilized by human participants.

## Notes

### Competing Interest Statement

The authors have declared no competing interest.

